# Physiological Significance of Bacterial Mn(II) Oxidation

**DOI:** 10.1101/2025.11.24.690301

**Authors:** Hui Lin, Yaohui Bai

## Abstract

Bacterial oxidation of Mn(II) to Mn(III/IV) oxides is widespread in nature and proceeds markedly faster than abiotic oxidation. However, the physiological role of this process remains unclear. Building on recent evidence that environmental stress determines bacterial Mn(II)-oxidation activity, we used *Pseudomonas putida* KT2440—a known Mn(II)-oxidizing bacterium—to investigate the potential drivers and benefits of this reaction. Using RNA-seq, we observed up-regulation of starvation-related two-component system genes during Mn(II) oxidation, supporting the idea that nutrient limitation acts as a trigger. This led us to hypothesize that introducing an external antagonistic stressor could further accelerate the onset and rate of Mn(II) oxidation. To test this, we established a synthetic microbial community in which KT2440 was confronted with *Pseudomonas aeruginosa* PAO1, which possesses both contact-dependent and diffusible antagonistic systems. Across multiple assays measuring oxidation initiation time, manganese oxide accumulation, and bacterial survival, the presence of PAO1 significantly accelerated Mn(II) oxidation by KT2440. More importantly, Mn(II) oxidation strongly enhanced KT2440 survival under antagonistic pressure. Correlation analysis indicated a positive relationship between survival rate and oxidation intensity. Fluorescence microscopy and agent-based modeling further confirmed that this protective effect is contact-dependent. By integrating these results with earlier findings, we propose that the physiological significance of bacterial Mn(II) oxidation lies in enhancing the survival fitness of the oxidizer under stressful conditions.

## Introduction

Manganese (Mn)—the second most abundant transition metal in Earth’s crust—underpins core biological functions, serving in superoxide dismutase and the oxygen-evolving complex of photosystem II (*1*). In natural systems, soluble Mn(II) is converted to insoluble Mn(III/IV) oxides via abiotic and biotic pathways; the latter proceed orders of magnitude faster (*2, 3*) and are mediated by diverse bacteria and fungi spanning the α-, β-, and γ-Proteobacteria, Actinobacteria, and Firmicutes across marine, freshwater, and terrestrial habitats (*4–10*). Microbial activity is therefore a principal driver of Mn oxides formation in environments ranging from metal-impacted streams to hydrothermal mounds and extensive ferromanganese deposits (*11–14*). Owing to their strong reactivity toward metals, organics, and oxidants, biogenic Mn oxides exert major control over redox transformations and the cycling of carbon and metals.

Despite this geochemical prominence, the cellular and ecological significance of Mn(II) oxidation remains unresolved (*15–18*). Proposed advantages include (i) energy conservation via electron transfer to Mn(III/IV) (*19, 20*), (ii) co-metabolic access to otherwise recalcitrant organic carbon through Mn-oxide-mediated transformations (*21, 22*) (iii) protection from reactive oxygen species and ultraviolet radiation, and (iv) detoxification via sorption of heavy metals and other toxins (*23*). Yet these putative benefits often lack decisive in vivo validation or are confounded by alternative defenses (e.g., catalases and superoxide dismutases) that can independently mitigate oxidative stress (*24, 25*). A promising route to clarify this puzzle is to ask when and why microbes activate Mn(II) oxidation. Recent work has implicated environmental stress and interspecies interactions as key triggers (*26*). In an *Arthrobacter* strain harboring a cryptic Mn(II) oxidation gene, exposure to stress—from either individual microbes or microbial consortia—can activate this silent pathway and initiate Mn(II) oxidation (*26*).

Here, we extend this framework to the tractable *Pseudomonas putida* KT2440, an individual Mn(II)-oxidizing bacterial strain. In monoculture with Mn(II), KT2440 begins to oxidize Mn(II) near the growth plateau and, at the same time, upregulates starvation-related two-component systems. These observations led us to hypothesize that a constant external antagonistic stress would (i) trigger earlier and faster Mn(II) oxidation and (ii) improve KT2440 survival during attack through this biomineralization. To test this, we built a synthetic attacker–target community in which *P. aeruginosa* PAO1, carrying both contact-dependent and diffusible attack systems, challenged Mn(II)-oxidizing KT2440 (*27, 28*). We measured when oxidation started, how fast it proceeded, how KT2440 populations changed under competition, and how cells and Mn oxides were spatially arranged. We employed agent-based modeling to reproduce these experimental patterns, which validated the contact-dependent nature of the process. Overall, we demonstrated bacterial Mn(II) oxidation as a stress-activated defensive strategy that enhances survival under competitive pressure by coupling mineral formation to heighten stress sensitivity.

## Materials and Methods

### Bacterial strains

Two *Pseudomonas* strains were used in the study, *P. aeruginosa* PAO1 (attacker, accession no. NBRC 100989) and *P. putida* KT2440 (Mn(II)-oxidizing bacteria, ATCC 11996). To visually distinguish the strains, plasmids expressing green fluorescent protein (GFP) or red fluorescent protein (RFP) mCherry (mCh) were introduced into PAO1 and KT2440, respectively, resulted in four strain variations: PAO1-GFP, PAO1-mCh, KT2440-GFP, and KT2440-mCh. All fluorescent proteins were expressed under the control of the arabinose-inducible promoter pRK found in plasmid pRK415. The fluorescent protein genes were incorporated into the plasmids, resulting in pRK415-EGFP and pRK415-mCherry. To ensure the stability of these plasmids within the bacterial cells, each plasmid conferred tetracycline antibiotic resistance to its host. Our observations revealed that while GFP-expressing strains grew at rates comparable to their non-fluorescent counterparts, the mCherry-expressing strains exhibited significantly reduced growth rates. These differences enabled us to simulate various ecological scenarios by altering fluorescent protein assignment to the strains. All bacterial strains, gene mutants, and plasmids used in this study are listed in Table S1.

### RNA sequencing

KT2440 strain were initially cultured in the modified PYG medium (*29*) (500 mg/L of MgSO_4_ ·7H_2_O, 60 mg/L of CaCl_2_ ·2H_2_O and 0.25 g/L of peptone, yeast extract and glucose) buffered by 10 mM 4-(2-hydroxyethyl)-1-piperazineethanesulfonic acid (HEPES) (pH 7.2), at 30°C with shaking at 170 rpm overnight. We sampled KT2440 suspensions before and after oxidation, at 6 h (exponential phase, no detectable Mn oxides, n=3) and 72 h (stationary-phase–like, strong Mn oxidation, n=3). Cell pellets were harvested by centrifugation at 10,000 *g* for 5 min at 4°C and immediately subjected to RNA extraction using TRNzol Reagent (TIANGEN, Beijing, China) according to the manufacturer’s instructions. RNA integrity was evaluated using an Agilent 2100 Bioanalyzer (Agilent Technologies, Santa Clara, CA, USA), and only samples with RNA integrity number (RIN) or RNA quality number (RQN) values greater than 8.0 were selected for cDNA library construction and sequencing. Libraries were sequenced on the Illumina NovaSeq 6000 platform, generating 250-bp paired-end reads. Raw sequence reads were processed to remove adapter sequences and low-quality reads using fastp (*30*), then ribosomal RNA were removed by mapped to the Rfam database (v14.9) using Bowtie2 (*31*). Clean reads were mapped to the reference genome sequences using Bowtie2 (*31*). Mapped reads were then counted using featureCounts (*32*) and subsequently analyzed for differential expression using the DEseq2 (*33*) algorithm in R (v4.4.1). Genes with an adjusted *P* value (Benjamini–Hochberg correction) ≤ 0.05 and |log₂ fold change| ≥ 1 were considered differentially expressed. KEGG-based functional annotation and pathway enrichment were then performed, with a focus on significantly upregulated genes assigned to the “two-component system” pathway to identify regulatory components potentially associated with Mn(II) oxidation.

### Construction of bacterial interaction system

We established a two-member community comprising the attacker *P. aeruginosa* PAO1 and the sensitive *P. putida* KT2440. PAO1 was chosen for its diverse arsenal of long-and short-range weapons. For long-range offense, PAO1 deploys phage-derived R-type pyocins (tailocins), which are particularly effective under biofilm-like conditions (*34, 35*). Although *P. aeruginosa* encodes a type VI secretion system (T6SS) for contact-dependent antagonism (*28*), its activation is tightly regulated and typically triggered only by direct threats (*36–38*). We therefore focused on PAO1’s contact-dependent inhibition (CDI) system as the principal short-range weapon under control conditions. The fitness costs of these weapons differ: tailocin production is SOS-induced and entails lysis of a subpopulation to release particles (*39*), whereas CDI, regulated by as-yet-unclear signals, can be produced without cell death (*40*). During growth, ∼3% of PAO1 cells lyse, releasing pyocins that diffuse through the extracellular milieu (diffusion coefficient on the order of 10⁻⁷ cm² s⁻¹) (*41, 42*) and can be taken up by sensitive KT2440 cells. Pyocin endonuclease activity degrades KT2440 DNA, halting division and culminating in lysis, with inhibition zones extending ∼100–400 μm from toxin producers (*43, 44*). By contrast, KT2440 lacks comparable antagonistic systems and grows markedly more slowly than PAO1 across inoculum densities, placing it at a clear disadvantage for limiting resources. We therefore used this defined two-member community as a controlled testbed to determine whether interspecies competition precipitates Mn(II) oxidation. Because spatial structure modulates both contact-dependent and diffusible antagonism, all assays were performed on agar to emulate dense, structured microenvironments typical of bacterial communities and to enable direct observation of competitive dynamics.

### Bacterial interaction assays

Overnight liquid precultures for KT2440 and PAO1 strain (both fluorescent and non-fluorescent) were independently cultured in LB medium, which contains 10 g of tryptone, 10 g of NaCl, and 5 g of yeast extract per liter, at 30 °C with agitation at 170 rpm. For the fluorescent strains, the medium was supplemented with 0.2% glycerol, 0.2% casein hydrolysate, and 0.2% arabinose to induce fluorescence, along with 20 µg/mL tetracycline to maintain plasmid stability. To assess the growth and survival of KT2440 strains (KT2440_mCh_ and KT2440_gfp_) in competition with PAO1 strains (PAO1 _gfp_ and PAO1 _mCh_), both with and without MnCl_2_ supplementation (0.1 mM), the overnight monocultures were washed twice with phosphate-buffered saline (PBS) and mixed at a 1:1 ratio. These mixtures were then serially diluted 10-fold (∼10^5^ to 10^9^), and 10-µL aliquots were spotted onto 1.5% w/v *Pseudomonas* agar medium (PDP) agar and incubated at 30 °C (Table S2). Once the cells began to grow on agar, bacterial warfare was initiated. This setup facilitated competition at various initial ratios and densities, with initial culture densities determined by serial dilution and spot plating.

### Measurement of Mn(II) Oxidation Rates

To quantify the Mn(II) oxidation rates in monoculture of KT2440 (KT2440_mCh_ and KT2440_gfp_) and co-cultures with PAO1 (PAO1 _gfp_ and PAO1 _mCh_), we utilized Inductively Coupled Plasma Optical Emission Spectrometry (ICP-OES, Shimadzu, Japan). Measurements were taken at various inoculum densities, ranging from approximately 10^5^ to 10^9^ cells, to assess the residual concentrations of Mn(II) in the medium. Samples were collected at 12 h intervals during the experiment to monitor the rate of Mn(II) conversion to Mn(III) or Mn(IV) oxides. The oxidation rates were calculated and expressed in milligrams of Mn(II) oxidized per liter per hour, providing a clear metric of the bacterial capability to oxidize manganese under various experimental conditions. This approach allows for precise determination of the bacterial oxidation activity under controlled experimental conditions.

### Imaging of colonies and quantification of competition outcomes

After 12 h of growth, images of colony formation were acquired using a Leica DMi8 inverted microscope (Leica THUNDER Imager Live Cell). Fluorescence was analyzed with specific filter sets: 472/30 nm excitation for GFP (DM: 495, BA: 520/35 BP) and 562/25 nm excitation for mCherry (DM: 593, BA: 641/45 BP). Image analysis was performed using ImageJ software (v2.1.0/1.53c). Strains expressing GFP appeared green, while those expressing RFP mCherry appeared red. This differentiation allowed for the identification of strains with a significant decrease in growth rate due to mCherry expression. Colonies from each group of frequencies and densities were photographed using the same magnification, ranging from 1× to 2.5×. Post-imaging, samples were collected from both the center and edge of each colony using a 10-µL pipette and suspended in PBS, followed by homogenization, serial dilution, and incubation at 30°C overnight. Colony counts were performed to establish the final ratio between the two strains. For quantifying colony-forming units (CFUs) of *Pseudomonas*, CN-agar plates (Hopebio cat# HB8484-2) supplemented with sulfamethoxazole (14.25 µg/mL), ampicillin (10 µg/mL), or tetracycline (10 µg/mL) were used to differentiate between the PAO1 and KT2440 strains (Table S3). CFU counting was performed using standard serial dilution and plating techniques. Approximately 10 µL of PBS-serially diluted cultures (dilution factors approximately 10^5^ to 10^9^) were spread onto 90-mm diameter agar plates using glass beads, with variations including the presence or absence of antibiotic selection.

### Agent-based model description

All agent-based simulations were performed in CellModeller (45), an open-source platform for modeling bacterial population dynamics. Additional Python modules were incorporated to implement contact-dependent and diffusible toxin secretion (46, 47); the corresponding source code is available on GitHub (https://github.com/WilliamPJSmith/CellModeller). To compare different antagonistic strategies, weapons were grouped into two classes—contact-dependent and diffusible—and their production was described by a common “secretion rate” parameter with normalized growth costs, allowing direct comparison across weapon types. Key processes, variables, and parameters used in the model are summarized in Table S4.

In the simulations, bacterial cells were modeled as rigid elastic rods that grow exponentially with a fixed radius *R*. They expand by pushing against each other as they elongate and divide once their initial volume *V*_0_ has doubled, incorporating a small random noise term ξ*V* ≈ *U*(0,0.05)*V*_0_. All cells had equal access to growth-limiting nutrients. During each simulation time step *Δ*t, the volume of each cell increased proportionally to its current volume, *V*’ = *k*_grow_*V*, discretized as *V*(*t* + Δ*t*) = *V*(*t*)(1 + *k*_grow_Δ*t*), where *k*_grow_(s^−1^) represents the net per capita growth rate. Weapon production, whether through diffusive or contact mechanisms, resulted in a growth penalty directly tied to the secretion rate, described by the formula *k*grow = *k*_max_(1 − *ck*_sec_). In this equation, *c* represents the proportional weapon cost, *k*_max_ is the maximum growth rate, and *k*_sec_ denotes the rate of contact weapon secretion. Following each growth cycle, an elastic energy minimization algorithm recalculated cell positioning to reduce overlap, also considering viscous drag on each cell. This method, thoroughly detailed in prior studies (*45–47*), effectively models the repulsive forces between physically contacting cells.

Simulations were initialized by randomly placing cells within a circular “homeland” of radius 50 µm on a flat surface. We systematically varied initial total cell density and the relative frequency of each strain. At time *t* = 0, cells were randomly positioned and oriented within this homeland. Each simulation was run for 100 time steps with Δ*t* = 0.025 h, and population dynamics and spatial organization were recorded over the course of the simulations.

## Results

### Starvation-induced Mn(II) oxidation

*P. putida* KT2440 is a Gammaproteobacterium and not a classical model Mn(II) oxidizer, but its close relatives (*P. putida* GB-1/MnB1) are well-established Mn(II)-oxidizing model strains (*48*). Genomic analysis of KT2440 has identified multicopper-oxidase–like genes homologous to the characterized Mn(II)-oxidizing enzymes in these relatives, indicating a latent capacity for biogenic Mn(II) oxidation (*49–51*). When grown with 0.1 mM MnCl₂, KT2440 began to produce Mn oxides as cultures approached the growth plateau at 24 h. Over this period, OD₆₀₀ increased from 0.050 ± 0.002 to a stable plateau of 1.766 ± 0.088, with no further exponential growth thereafter (Fig. 1a). Over the same interval, dissolved Mn(II) decreased from 6.21 mg·L⁻¹ to 2.29 mg·L⁻¹ at 24 h (Δ = 63.2% reduction), consistent with pseudo–first-order loss kinetics (fit: kₒₓ = 0.069 h⁻¹, R² = 0.90; Fig. 1a).

**Figure 1.**
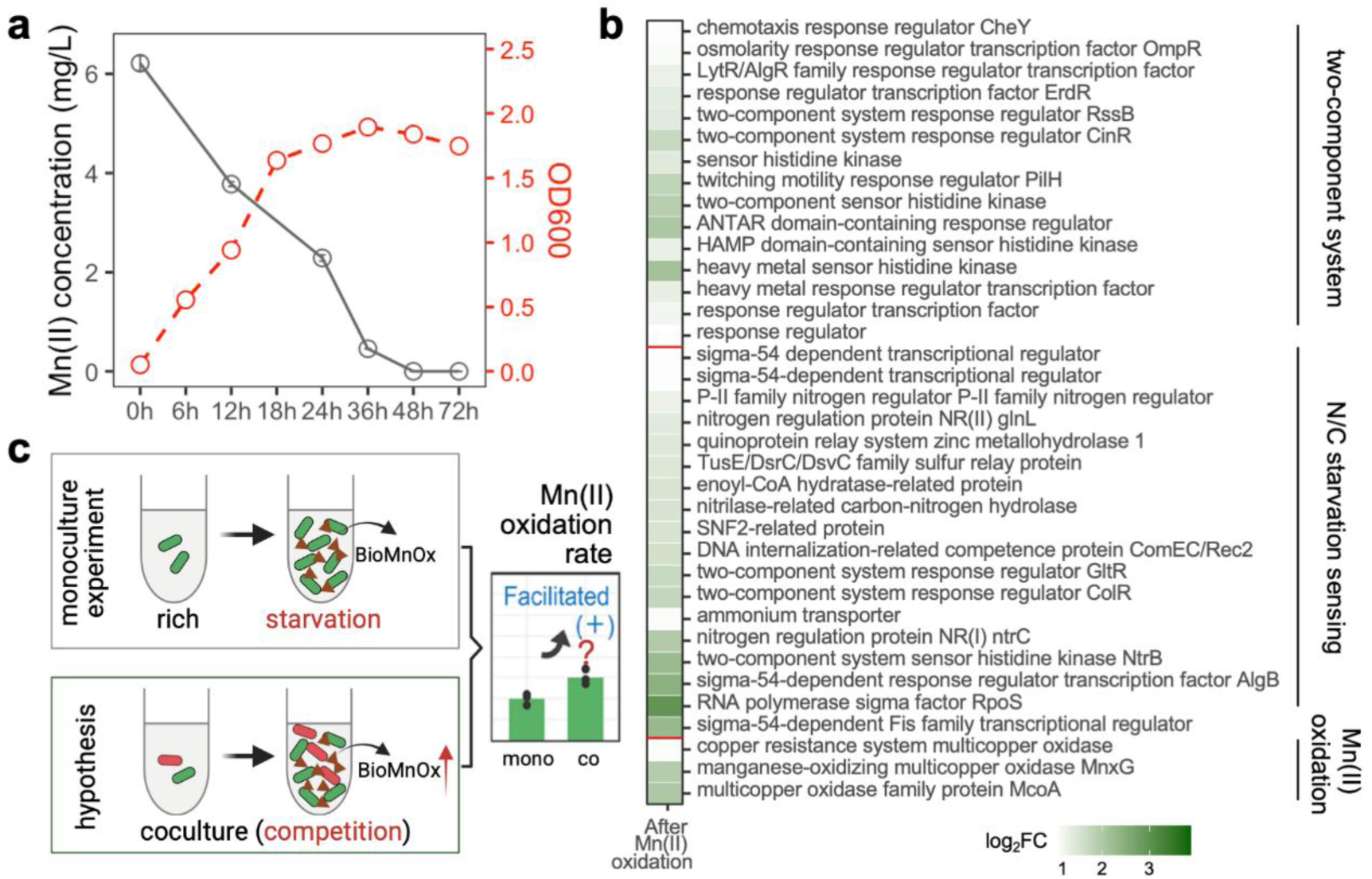
Mn(II) oxidation in KT2440 and stress-linked transcriptional response. (**a**) Growth (OD₆₀₀) and dissolved Mn(II) in *P. putida* KT2440 monocultures supplied with 0.1 mM MnCl₂. (**b**) Differential gene expression at 6 h (exponential phase, no Mn oxides, n=3) versus 72 h (stationary-phase–like, strong Mn oxidation n=3), showing enrichment of two-component systems and starvation-sensing genes, and upregulation of multicopper-oxidase–like Mn(II) oxidation genes (*mcoA*, *mnxG*). (**c**) Proposed hypothesis: interspecies competition, by adding stress, is predicted to trigger earlier onset and faster Mn(II) oxidation.

To link Mn(II) oxidation to cellular physiology, we performed RNA-seq on KT2440 cells sampled before and after oxidation, at 6 h (exponential phase, no detectable Mn oxides) and 72 h (stationary-phase–like, strong Mn oxidation). Guided by our previous work on stress-activated Mn oxidation, we focused on differentially expressed genes in two-component regulatory systems. Two-component signal transduction genes were significantly enriched (NES = 1.8; [FDR] *q* < 0.05), consistent with activation of downstream starvation– and stress-response pathways, and putative Mn(II) oxidation machinery—multicopper-oxidase–like oxidoreductases—was upregulated at onset (e.g., *mcoA* and *mnxG*: log₂FC = 1.931 and 2.014; [FDR] *q* < 0.01, Fig. 1b), linking this stress state to activation of the oxidative pathway. Together, these data show that KT2440 engages Mn(II) oxidation after entering a nutrient-limited, stationary-phase–like state, rather than during exponential growth. This supports the view that Mn(II) oxidation is a stress-linked response and motivates the idea that interspecies competition—by intensifying nutrient scarcity and adding antagonistic pressure—could further advance the onset and increase the rate of Mn(II) oxidation (Fig. 1c).

### Interspecies competition accelerates Mn(II) oxidation

We spotted 10-µL cocultures, KT2440 and PAO1, spanning 10-fold inoculum dilutions (10^5^ to 10^9^ CFU/mL) onto agar with or without MnCl₂ and monitored colonies for 48 h. In the absence of MnCl₂, colonies expanded rapidly and developed increasingly serrated rims over time, consistent with edge instabilities under progressive nutrient depletion (*52, 53*). With MnCl₂, colonies were smaller, more circular, and exhibited smoother margins (Fig. 2a and Fig. S1).

**Figure 2.**
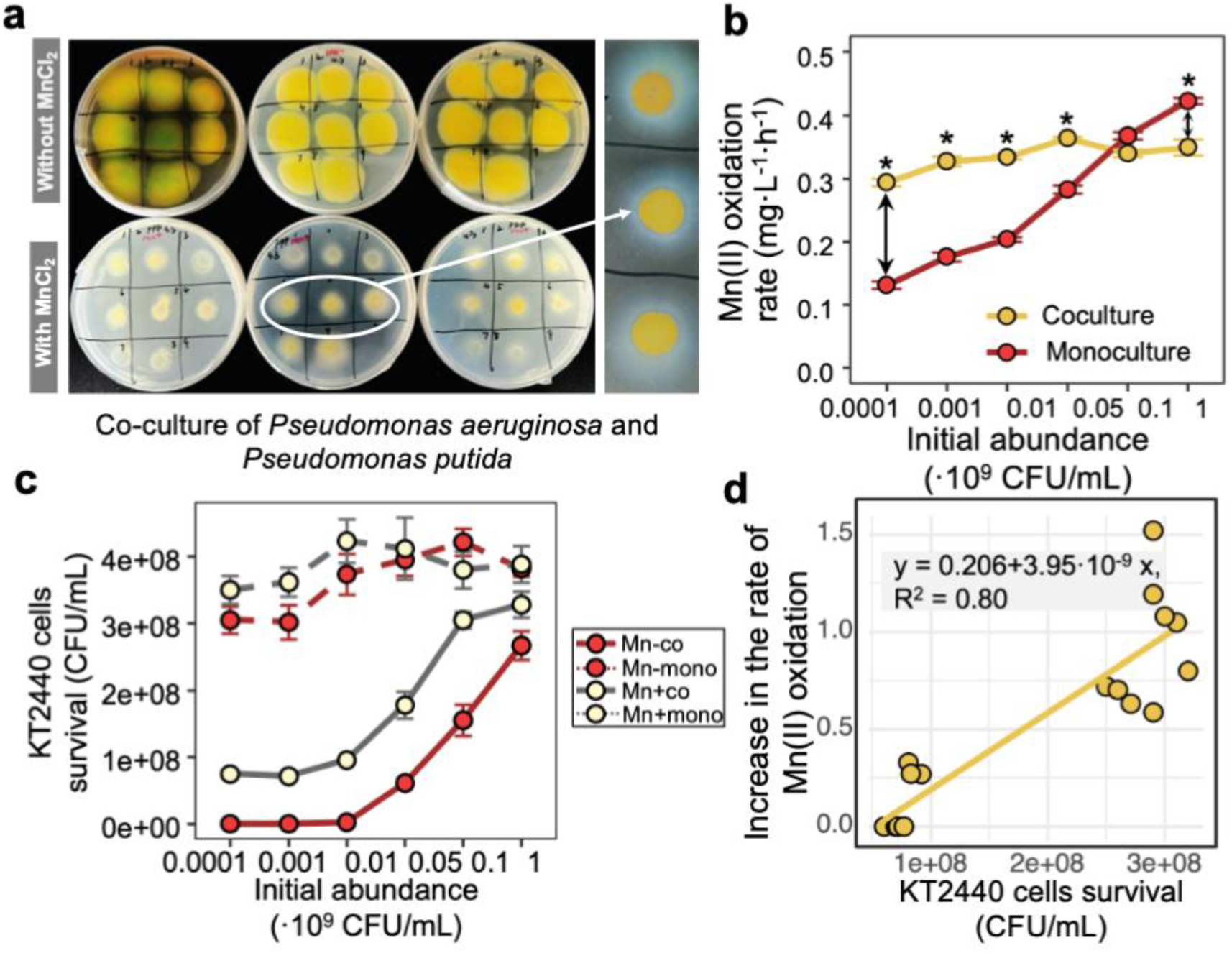
Growth dynamics of *Pseudomonas aeruginosa* PAO1 and *Pseudomonas putida* KT2440 co-cultures on agar plates. (**a**) Colonies without MnCl₂ exhibited rapid expansion and increased margin irregularity, highlighting dynamic interactions within the microbial community. (**b**) Mn(II) oxidation rates by KT2440, assessed across various inoculum densities in co-culture with PAO1 and in monoculture (n = 3), showed significant differences at lower densities. (**c**) Following MnCl₂ addition, KT2440 demonstrated significantly enhanced survival at lower inoculum densities (n = 6), accompanied by visible brown Mn oxides after 12 h, suggesting a protective role of these oxides. Significance (\**p* < 0.05) was assessed using the Wilcoxon test. (**d**) The relationship between survival and incremental Mn(II) oxidation rate in KT2440. Equation depicted the relationship, while R^2^ indicated the linear correlation between survival and Mn(II) oxidation rate. In all regression analyses, the T-test yielded *p*-values less than 0.05.

We first quantified Mn(II) oxidation by KT2440 in monoculture versus coculture. In monoculture, oxidation displayed strong density dependence, increasing markedly with higher starting densities. Under co-culture conditions, although Mn(II) oxidation rates remained similar across different inoculum densities, they exhibited an inoculum-dependent effect compared to the rate increase observed in single cultures. At lower inoculum densities (ranging from 10^5^ to 10^7^ CFU/mL), Mn(II) oxidation significantly increased under co-culture conditions compared to monoculture conditions (Fig. 2b and Fig. S2). Conversely, at higher inoculum densities (10^8^ CFU/mL), there was no significant difference in Mn(II) oxidation rates, and in some instances (10^9^ CFU/mL), Mn(II) oxidation was suppressed (Fig. 2b and Fig. S2). Thus, competition elicits an early, density-sensitive acceleration of Mn(II) oxidation that dissipates—and can even invert—at very high starting densities.

We next quantified competitive outcomes by selective plating. Enabling Mn-oxide formation with MnCl₂ significantly improved KT2440 survival at low inocula (ranging from 10^5^ to 10^7^ CFU/mL) in coculture (Fig. 2b and Fig. S3). At the lowest inoculum (∼10^5^ CFU/mL), removing MnCl₂ caused an ∼10^6^-fold loss of KT2440 (Fig. 2c and Fig. S3), consistent with PAO1’s advantage via both contact-dependent inhibition (CDI) and diffusible pyocins; conversely, MnCl₂ markedly mitigated this loss, underscoring the protective role of Mn oxides (Fig. 2c). Additionally, MnCl₂ did not measurably affect KT2440 growth rate or biomass across starting densities (Fig. S4), indicating a defensive benefit without detectable growth cost.

KT2440 survival was significantly correlated with the density-dependent acceleration of Mn(II) oxidation (Fig. 2d). Transmission electron microscopy (TEM) revealed a conspicuous pericellular Mn-oxide ring around KT2440 (Fig. S5), effectively encapsulating them and creating an inhibition zone. To further differentiate the protective effects of Mn oxides against both short– and long-range weapons, we measured pyocin production by PAO1. While the addition of MnCl_2_ significantly reduced pyocin production (Fig. S6a), the introduction of sterile Mn oxides did not significantly degrade pyocins (Fig. S6b). These findings suggest that Mn oxides may primarily offer protection against short-range contact weapons, rather than degrading chemical toxins.

### Mn(II) oxidation increases bacterial survival fitness

To investigate whether Mn(II) oxidation enhances KT2440’s fitness (survival), we fluorescently labeled the strains by engineering *P. aeruginosa* PAO1 and *P. putida* KT2440 to express GFP and mCherry, respectively. Observations indicated that the PAO1_gfp_ strain expanded at a rate comparable to its non-fluorescent counterpart, while KT2440_mCh_ showed a significant 22.6% reduction in growth (Fig. S7a). However, the relative growth rates between the two fluorescent strains remained consistent across different initial cell densities compared to the non-fluorescent strain (Fig. S7a). Consistent with this, Mn(II) oxidation dynamics in coculture remained similar to the unlabeled system: at low inoculum densities, the oxidation rate increased with decreasing density and was ∼3-fold higher at an initial density of 10⁵ CFU/mL (Fig. S7b).

Competition experiments using these fluorescent protein-expressing strains—PAO1_gfp_ (green) and KT2440_mCh_ (red)—were conducted with and without MnCl_2_ supplementation on standard agar plates. Images of the co-cultures were captured to analyze the spatial distribution of the attacking and susceptible MnOB. In the absence of MnCl_2_, KT2440_mCh_ cells were predominantly single and located at the colony edge (Fig. 3a). In the presence of MnCl_2_, at lower densities (< 10^7^ CFU/mL), the bacteria were encapsulated in Mn oxide patches scattered across the colony center, enhancing structural integrity and survival (Fig. 3b). At higher densities (> 10^7^ CFU/mL), these formations aggregated towards the colony edge (Fig. 3b). These fluorescent patches initially appeared after 12-h of co-culture during the early stages of the Mn(II) oxidation process (Fig. S8a) and became more pronounced when brown Mn oxides were visible to the naked eye on agar plate by 24-h of co-culture (Figs. S8b and S9a). After an extended co-culture period of 96-h, the centers of the plate colonies turned a completely dark brown color, regardless of the initial inoculum density (Fig. S9b-c). Quantitative analysis following colony elution and resuspension confirmed that the presence of Mn oxides significantly enhanced KT2440_mCh_ survival across various densities (Fig. 3c).

**Figure 3.**
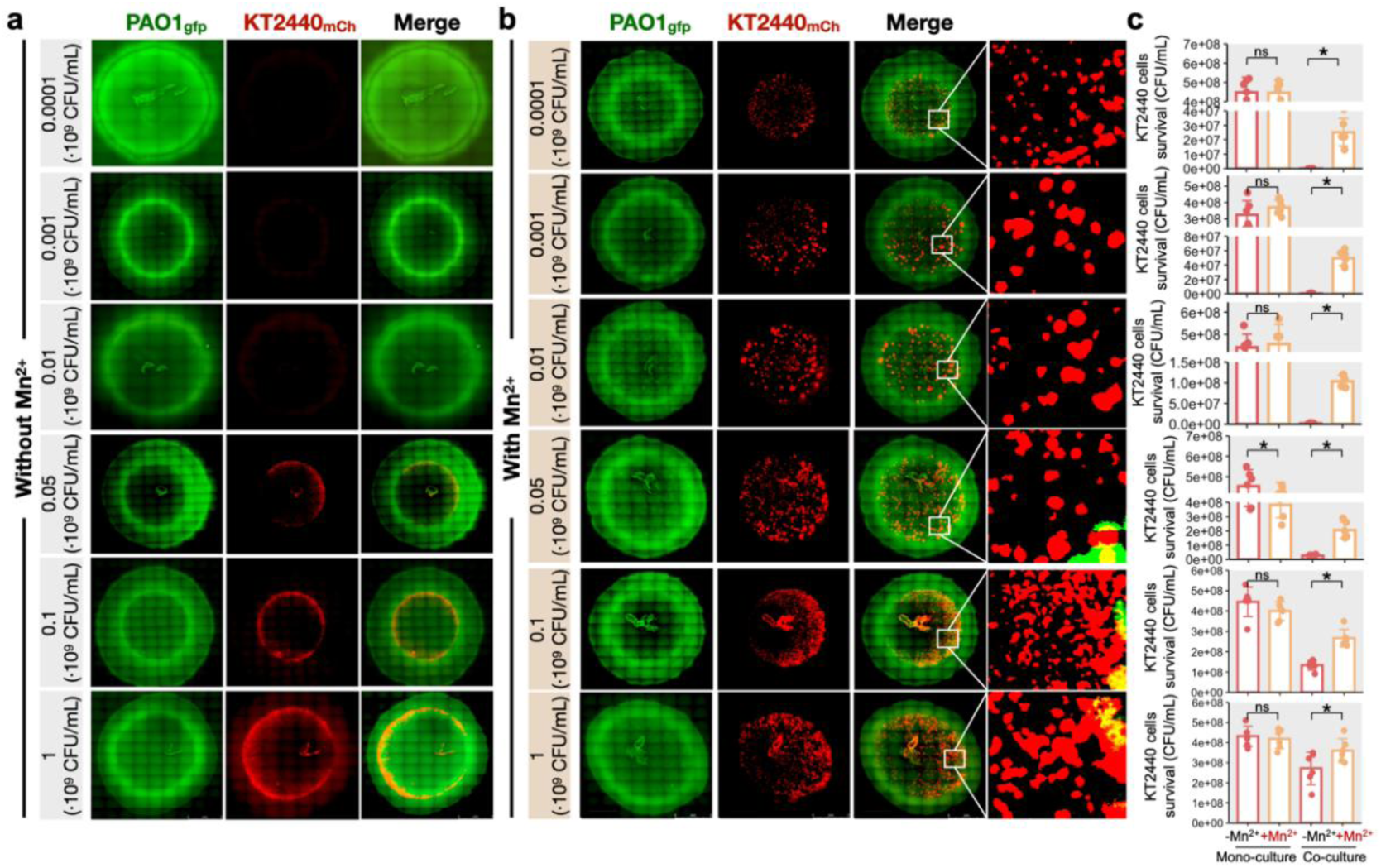
Fluorescence microscopy of the *Pseudomonas aeruginosa* PAO1_gfp_ and *Pseudomonas putida* KT2440_mCh_ co-cultures on agar plates. We co-cultured PAO1 (expressing GFP) and KT2440 (expressing mCherry, RFP) cells: (**a**) without MnCl₂ and (**b**) with MnCl₂ after 24 h of growth. In the presence of MnCl₂, especially at lower densities, KT2440_mCh_ cells were encapsulated in Mn oxide patches distributed throughout the colony center. (**c**) This encapsulation led to a significant increase in KT2440_mCh_ survival across various densities (n = 6). Statistical significance (\**p* < 0.05) was determined using the Wilcoxon test.

We next examined a low-competition regime by inverting growth-rate ranks between the attacker (PAO1) and the susceptible MnOB (KT2440). The fluorescent proteins were swapped between the two strains, with PAO1 expressing mCherry and KT2440 expressing GFP. This modification was based on earlier findings that mCherry-expressing strains typically exhibited reduced growth rates compared to wild-type strains, unlike those expressing GFP. Specifically, KT2440 tagged with GFP (KT2440_gfp_) demonstrated growth similar to its wild-type counterpart, whereas PAO1 expressing mCherry (PAO1_mCh_) showed a notable 36.7% ± 10.8% decrease in growth compared to wild-type cells across initial cell densities ranging from 10^5^ to 10^9^ CFU/mL (Fig. S10a). Fluorescence micrographs demonstrated that at varying initial densities, the presence of Mn oxides led to patchy growth patterns in KT2440_gfp_ cells (green), similar to observations before the reversal of growth rate classifications (Fig. 5a-b). Despite these patterns, there was no significant impact on cell density in the co-cultures and monocultures of KT2440_gfp_ or PAO1_mCh_, regardless of MnCl_2_ addition (Fig. 5c). Furthermore, Mn(II) oxidation rates did not differ between co-cultures and monocultures (Fig. S10b), indicating that when interspecific antagonism is weak—i.e., the susceptible strain is not growth-disadvantaged—oxidation is not elevated. Consequently, Mn oxides neither improve survival nor shift competitive outcomes under these conditions.

**Figure 4.**
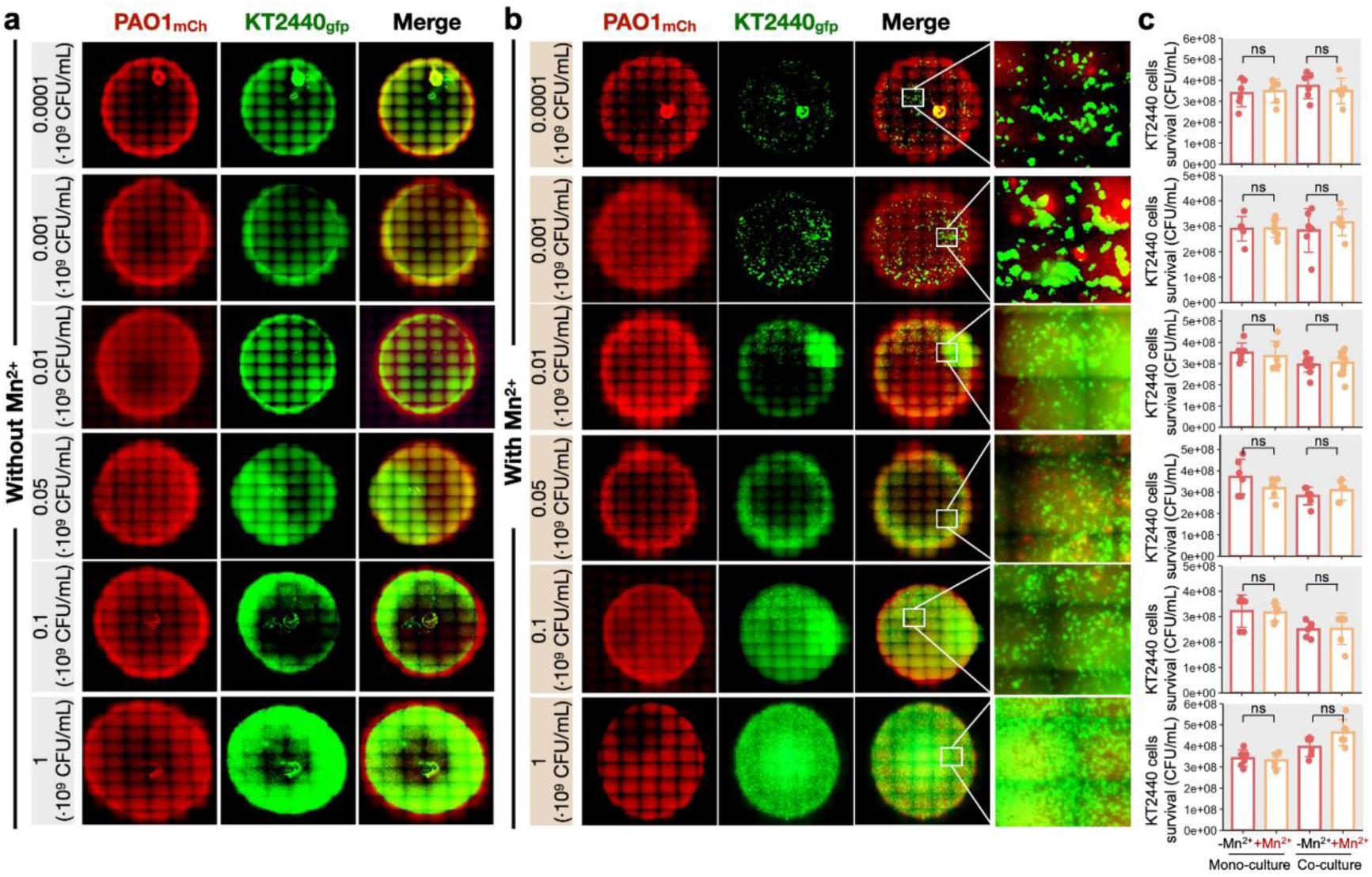
Competitive dynamics between *Pseudomonas aeruginosa* PAO1_mCh_ and *Pseudomonas putida* KT2440_gfp_ with reversed growth rate rankings. PAO1 (expressing mCherry) and KT2440 (expressing GFP) were co-cultured on agar plates to assess competitive interactions (**a**) without MnCl₂ and (**b**) with MnCl₂ after 24 h of growth. (**c**) Mn oxides did not significantly impact the overall composition of the competitive community (n = 6). Statistical significance (\**p* < 0.05) was determined using Wilcoxon test, indicating minimal impact on competitive outcomes despite reversed growth rate rankings.

**Figure 5.**
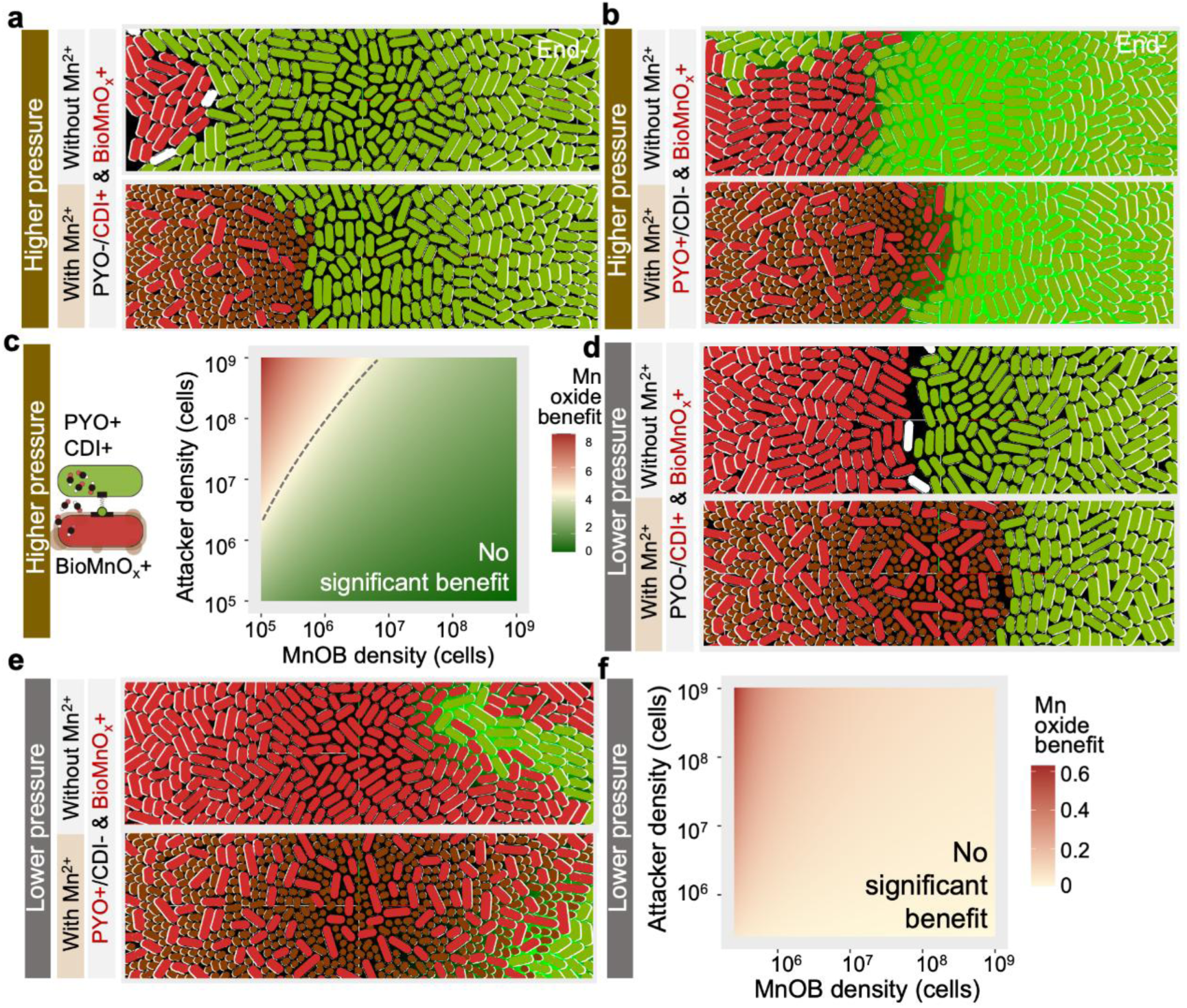
Agent-based model of competition between bacteria with varied weaponry and susceptible Mn(II)-oxidizing bacteria (MnOB). (**a**) Simulations with attackers bearing only short-range (contact-dependent) weapons. (**b**) Simulations with attackers bearing only long-range (diffusible) weapons. (**c**) Protective efficacy is density-dependent—strongest when attacker inoculum is high and MnOB inoculum is low. (**d**)-(**f**) Scenarios with reversed growth-rate ranking (attacker ≤ MnOB), evaluated for short-range (**d**) and long-range (**e**) attackers, to probe low-competition regimes.

### Agent-based modeling suggests Mn(II) oxidation was contact-dependent

To determine which antagonistic mechanisms trigger protective Mn(II) oxidation, we used an agent-based model of competition between an attacker and a susceptible Mn(II) oxidation strain (MnOB). Across a wide range of parameter combinations (weapon type, toxin secretion rate, and Mn-oxide secretion), Mn oxides consistently provided strong protection when attackers relied on contact-dependent weapons, but offered little benefit against purely diffusible toxins (Fig. 5a-c, Movie S1). Next, we conducted parameter sweeps in the simulation to determine the conditions under which Mn oxide production offers protection against both short– and long-range attacks. By contrasting simulations featuring Mn oxides (BioMnO_x_+) with those without (BioMnO_x_−, at rates of 0.1 and 0 respectively), we employed a straightforward metric to evaluate the protective advantages of Mn oxide production:

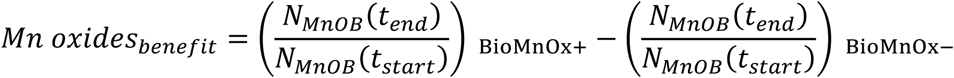

where *N*_*MnOB*_(t) is the cell density of surviving MnOB present at time t. Therefore, the benefits of Mn oxides are evident in the survival rates of Mn(II)-oxidizing bacteria, relative to their initial populations, compared to those that do not produce oxides. Simulations with varying numbers of attackers and susceptible MnOB (using the same *N*_*MnOB*_(*t*_*start*_) for BioMnO_x_+ and BioMnO_x_− simulations) consistently showed that Mn oxide production provided at least some protection against both short and long-range weapons (Mn oxides benefit > 0). This protective effect was most pronounced under a higher density of attackers compared to a lower density of susceptible MnOB (Fig. 5d). In simulations mimicking invasion into an established MnOB biofilm, increased Mn-oxide secretion markedly reduced the success of short-range attackers, but only weakly impeded long-range attackers, which eventually invaded given sufficient time (Fig. S11). Finally, when we relaxed competitive pressure by reversing the growth-rate ranking of the two strains, Mn oxides no longer improved MnOB survival, irrespective of whether attackers used short– or long-range weapons (Fig. 5d-f). Together, these simulations support a simple conclusion: loose, non-membrane-bound Mn oxides primarily function as a defensive barrier against contact-dependent killing under strong interspecific competition, with limited efficacy against diffusible toxins or under weak antagonism.

## Discussion

A fundamental question in bacterial Mn(II) oxidation remains unresolved: the motive for this oxidation process (*2, 54*). Some studies propose that Mn(II) oxidation is an incidental byproduct of nonspecific reactions with cellular or extracellular compounds (*54*), while others suggest it may be an evolutionary relic with little current physiological relevance (*55*). However, the prevalence of Mn(II) oxidation among diverse bacteria (*2, 56, 57*), and the tendency of many Mn(II)-oxidizing bacteria to become encrusted in Mn oxides (*2, 58–60*), implies that there may be significant evolutionary advantages to possessing this trait. Hypothesis is that Mn oxides provide protection from various environmental insults, such as UV radiation, predation, viral attack, and heavy metal toxicity (*61–63*). However, decisive *in vivo* evidence supporting these protective roles has remained scarce.

Our results provide direct evidence that bacterial Mn(II) oxidation can function as a stress-activated survival mechanism. In *P. putida* KT2440, Mn(II) oxidation in monoculture begins only late in growth, when starvation-related signals emerge. When an antagonist (*P. aeruginosa* PAO1) is present, KT2440 initiates Mn(II) oxidation earlier and more strongly, showing that external competitive stress is integrated with internal nutrient stress to regulate this pathway. This pattern is consistent with our previous work in *Arthrobacter*, where a Mn(II)-oxidizing gene (*boxA*) remains silent in monoculture and is activated only when a stress-sensing system detects appropriate signals (*26*).

This perspective helps explain why Mn(II) oxidation is widely maintained across bacteria. A recent genomic survey found that roughly 23% of environmental bacterial genomes encode homologs of Mn(II)-oxidizing enzymes (*64*), yet many of these organisms do not produce Mn oxides under benign conditions. Our findings suggest that, in natural settings where microbes frequently encounter competition, predation, or fluctuating harsh conditions, a latent Mn-oxidation capacity functions as a physiological reserve. When stress is sensed (e.g. nutrient exhaustion, microbial antagonism, or exposure to oxidants), cells can rapidly activate Mn-oxidizing enzymes to generate biomineral Mn oxides that restructure their immediate microenvironment and support survival under challenge. The Mn oxides that precipitate around cells modify cell–cell contacts and local chemical gradients, effectively amplifying the impact of stress signals and locking the community into a more responsive state. Thus, bacteria oxidize Mn(II) to create a stress-induced mineral interface that enhances how acutely they perceive and respond to environmental challenges.

## Data availability

All code for the simulations was performed in CellModeller, available at https://github.com/cellmodeller/CellModeller. Additionally, transcriptome sequence data for *Pseudomonas putida* KT2440 has been deposited in the China National Center for Bioinformation database (https://www.cncb.ac.cn) under accession number CRA033685. Other datasets and code generated during and/or analysed during the current study were available from the corresponding author on reasonable request.

## Acknowledgements

This work was supported by the National Natural Science Foundation of China (52450009).

## Author contributions

H.L. conducted the experiments, performed data analyses, and prepared the figures.

H.L and Y.B. together designed the study and wrote the manuscript. Y.B. applied for funding.

