## Supplemental Materials for "Physiological Significance of Bacterial Mn(II) Oxidation"

**Competing interests**

The authors declare no competing interests.

**SUPPLEMENTARY VIDEO LEGEND**

**Movie S1.** Agent-based model of bacterial competition. This video simulation shows the interactions between bacteria armed with both short- and long-range weapons and susceptible Mn(II)-oxidizing bacteria (MnOB). It contrasts scenarios without and with Mn oxides, highlighting how Mn oxides influence the dynamics of microbial competition.

**SUPPLEMENTARY METHODS**

***Supplementary Method S1*** ***Pyocyanin accumulation in cultures***

Co-cultures of *Pseudomonas putida* KT2440 and *Pseudomonas aeruginosa* PAO1, both with and without MnCl₂ addition, were harvested at the exponential phase (24 h) to quantify excreted pyocyanin, which appears as a pink to deep red color in an acidic solution. From each 96-well plate containing PDP-agar co-cultures, 600 μL was mashed and mixed with 360 μL of chloroform to extract pyocyanin. The chloroform phase was then transferred to 120 μL of 0.2 M hydrochloric acid (HCl). After centrifugation at 5,000 ×g for 2 minutes, the absorbance of the top aqueous layer was measured at 520 nm. Pyocyanin concentrations were calculated as micrograms per milliliter of supernatant by multiplying the absorbance (A520 nm) by 17.1.

***Supplementary Method S2 Biological transmission electron microscopy (TEM)***

Co-cultures of *Pseudomonas putida* KT2440 and *Pseudomonas aeruginosa* PAO1 were fixed immediately after 24 h of growth and complete manganese oxidation using 2.5% glutaraldehyde, and stored at 4 °C for 12–24 h for primary fixation. Samples were then post-fixed in a solution of 1% osmium tetroxide and 2% potassium ferrocyanide for 1–2 h, followed by four rinses with 18.25 MΩ deionized water, 15 minutes each. The samples underwent dehydration through a graded series of acetone concentrations from 30% to 100%, increasing in 20% increments, each step lasting 10 minutes, with the final step at 100% acetone repeated twice for 20 minutes each. Embedding was performed in stages beginning with a mix of acetone and embedding medium at 3:1 and 1:1 ratios at 37 °C for 1 and 3 h, respectively, then in pure embedding medium at 37 °C overnight. Samples were polymerized in embedding molds at 60 °C for 48 h. Ultrathin sections (70–90 nm) were cut using a Leica UC7 ultramicrotome (Leica Microsystems, Germany), placed on copper grids, and stained with uranyl acetate for 15 minutes and lead citrate for 10 minutes. The stained sections were analyzed under a Hitachi HT7800 transmission electron microscope (TEM) (Hitachi High-Technologies Corporation, Japan) for ultrastructural examination.

***Supplementary Method S3 Agent-based model: Contact-dependent weapons***

Our simulation utilizes a Python module designed to model antagonistic interactions between cells via contact-dependent mechanisms (*1*), including T6SS-mediated interference and contact-dependent inhibition (CDI). In each simulation step, cells with contact toxins extended ‘needles’ of length 𝑅 orthogonally from random points on their surfaces. The occurrence rate of these secretion events per cell over time adhered to a Poisson distribution, with the average frequency determined by the secretion rate 𝑘_sec_. Following each needle discharge, the line-segment technique was used to ascertain if it contacted any other cell in the population. Hits on target cells were documented, and a cell was deemed deceased if the tally of hits it sustained (excluding those from kin cells) exceeded a lethal threshold, set at 𝑁_hits_ = 1 (*2*).

***Supplementary Method S4 Agent-based model: Diffusible-dependent toxins***

We modeled toxins as freely diffusible solutes that inhibit the growth of susceptible cells when their local concentration 𝑢_𝑇_ exceeds a lethal threshold 𝑇_𝐶_, following the formula: $MnOB.growthRate=\frac{V_{max}}{1+\frac{Attacker}{K_{m}}}$. To model the toxin concentration field 𝑢_𝑇_ = 𝑢_𝑇_(𝑥,𝑦) (kg_T_ m^−3^) for a given cell configuration, we used the reaction-diffusion equation: $\frac{\partial u_{T}}{\partial t}=D_{T}\nabla^{2}u_{T}+K_{T}\alpha\rho\phi(x,y)$, where 𝐷_𝑇_, 𝑘_𝑇_, 𝛼, 𝜌, and 𝜙(𝑥,𝑦) represent the toxin diffusivity (m^-3^), specific toxin production rate (s^-1^), toxin yield per unit cell biomass (kg_T_kg_X_^-1^), cell biomass density (kg_X_ m^-3^), and cell volume fraction function (unitless), respectively. In non-dimensional form, pseudo-steady-state solutions to this equation can be represented by: $\nabla^{2}u_{T}=D_{T}\emptyset(x,y)$, where $D_{T}$is the Damköhler number defined as $D_{T}=\frac{l^{2}K_{T}\alpha\rho}{D_{T}T_{C}}$. We derived pseudo-steady-state solutions using the finite element method, incorporating the relevant CellModeller signaling modules. These solutions were analyzed within a two-dimensional (2D) rectangular domain of dimensions 𝐿_𝑥_ by *L_y_*, subject to mixed boundary conditions: periodic on the left and right sides, Neumann at the bottom, and Dirichlet conditions (where 𝑢_𝑇_ = 0) at the top. It was presumed that toxin degradation or removal occurred only through leakage at the top boundary.

***Supplementary Method S5 Simulation protocol***

We implemented all simulations in an agent-based framework where each bacterial cell is represented as an individual agent on a 2D surface mimicking a biofilm or mucosal layer. Cells interacted mechanically by pushing against each other as they grew and collided, and chemically by secreting toxins that inhibited susceptible genotypes (Movie S1). When cells exceeded a prescribed height, they were removed from the simulation to represent dispersal or sloughing from the top of the community. In our simulations, attacker cells either ‘fired’ a short-range weapon randomly from their cell surface, impacting nearby susceptible cells to simulate contact-dependent systems such as T6SSs and CDIs, or released a diffusible substance, representing various toxins such as small-molecule antibiotics, ribosomally synthesized bacteriocins, or tailocins. The secretion rates of these toxins, whether from short- or long-range weapons, were standardized and regulated by the secretion rate parameter ksec. The lethality of the toxins was matched, with both contact and diffusible toxins becoming lethal when their intracellular concentrations exceeded the threshold Tc. For Mn(II)-oxidizing bacteria, Mn oxide production was modeled as the secretion of loose, spherical particles into the environment, reflecting their interaction with surrounding cells and medium.

**SUPPLEMENTARY FIGURES**

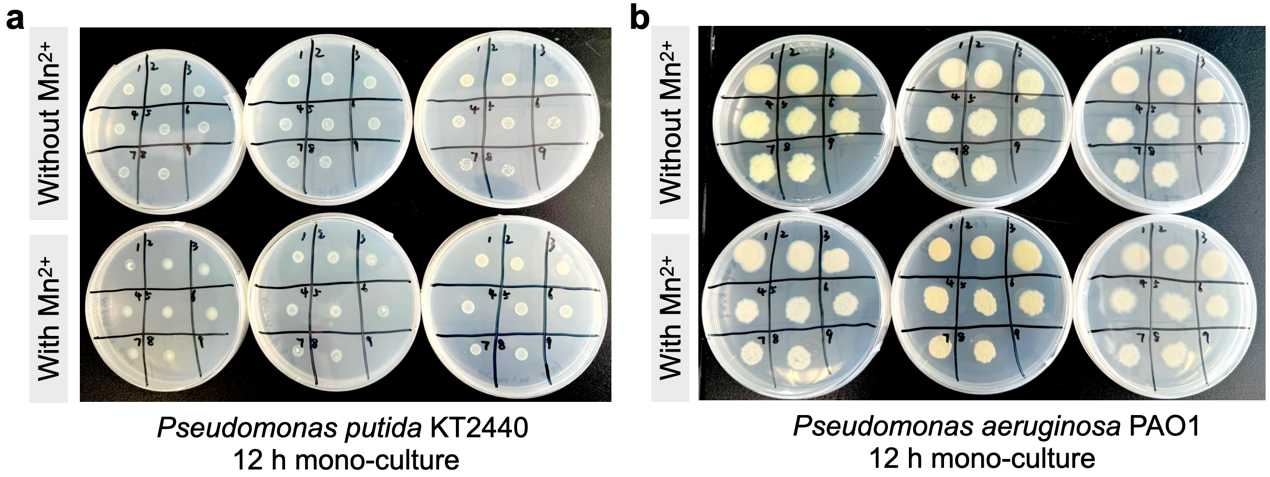

**Figure S1.** Growth patterns of *P. putida* KT2440 and *P. aeruginosa* PAO1 on agar plates after 12 h of culture. (**a**) KT2440 cells displayed vertical growth and maintained a circular shape. (**b**) PAO1 colonies spread horizontally, leading to larger and more irregular shapes.

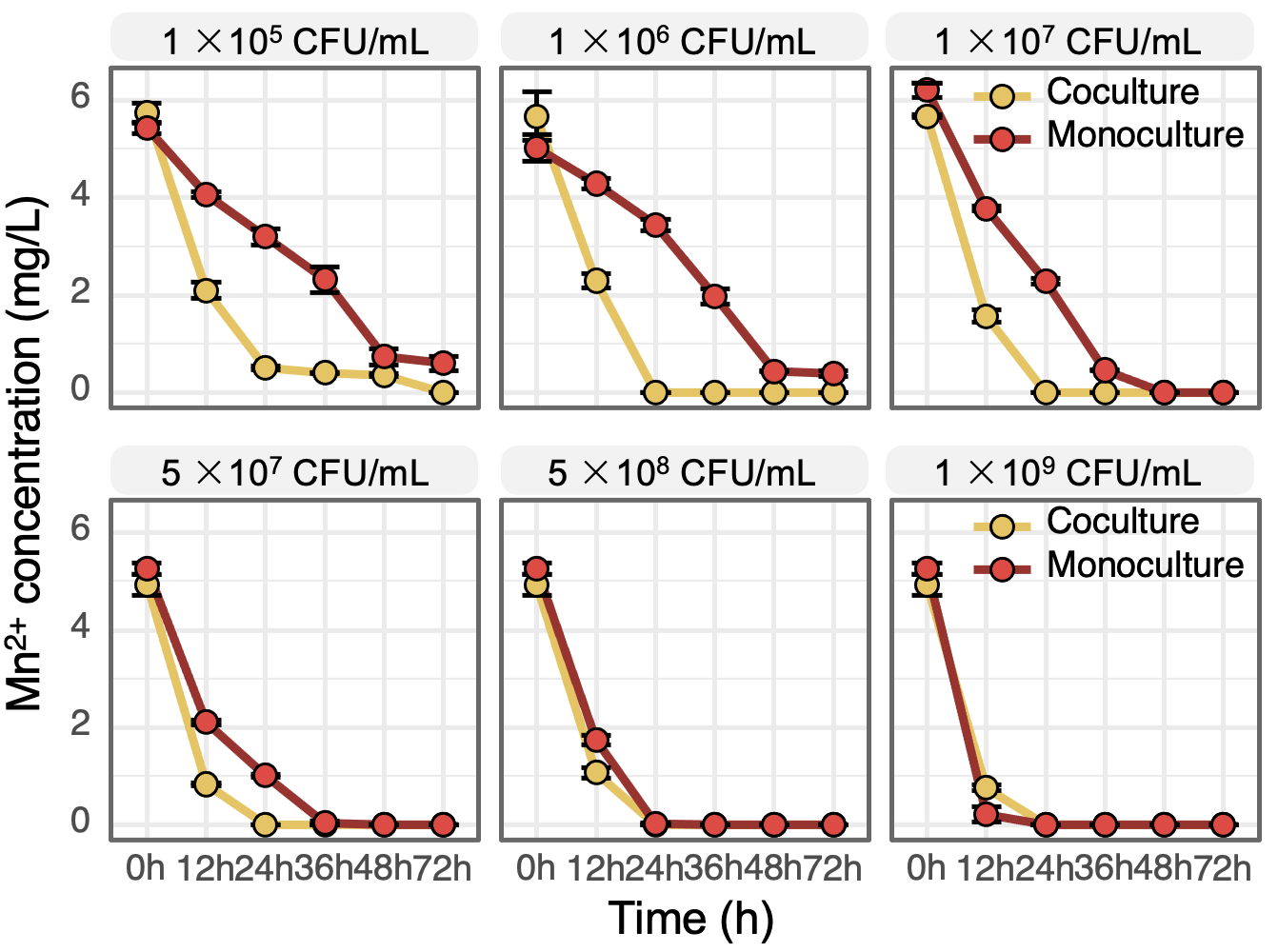

**Figure S2.** Changes in Mn^2+^ concentration over time under *P. putida* KT2440 monoculture or co-culture with *P. aeruginosa* PAO1 (n = 3).

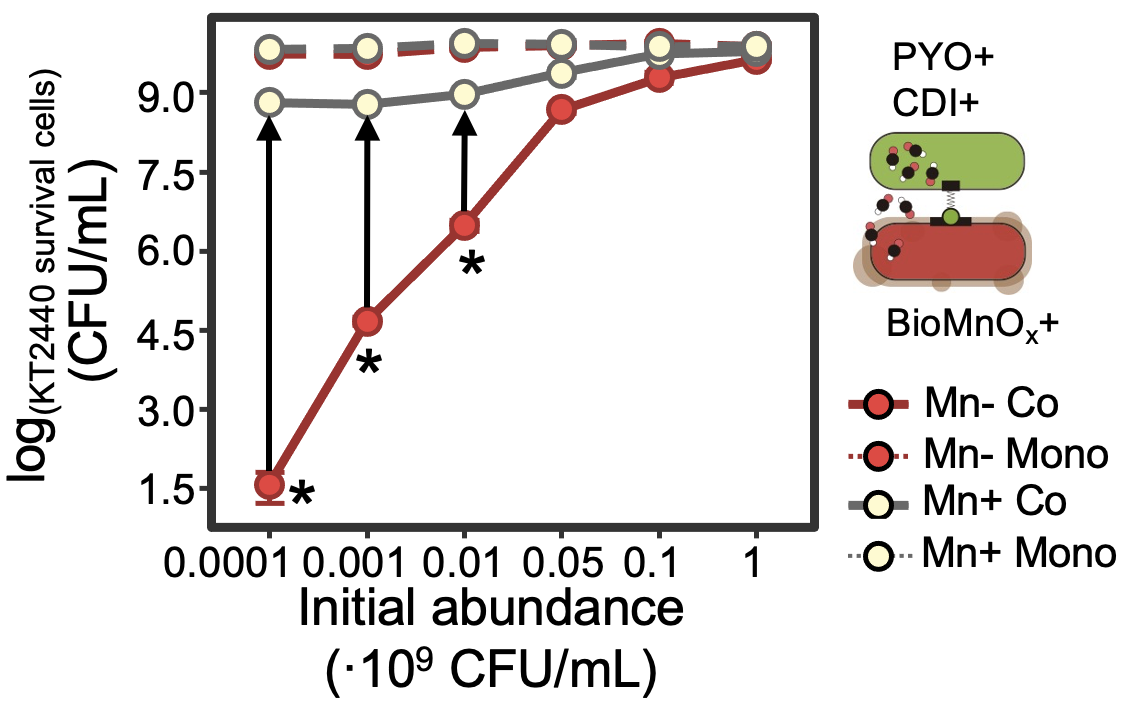

**Figure S3.** Log-transfer survival of *P. putida* KT2440 across varying initial inoculum densities in monoculture and co-culture with *P. aeruginosa* PAO1 with and without MnCl₂ (n = 6).

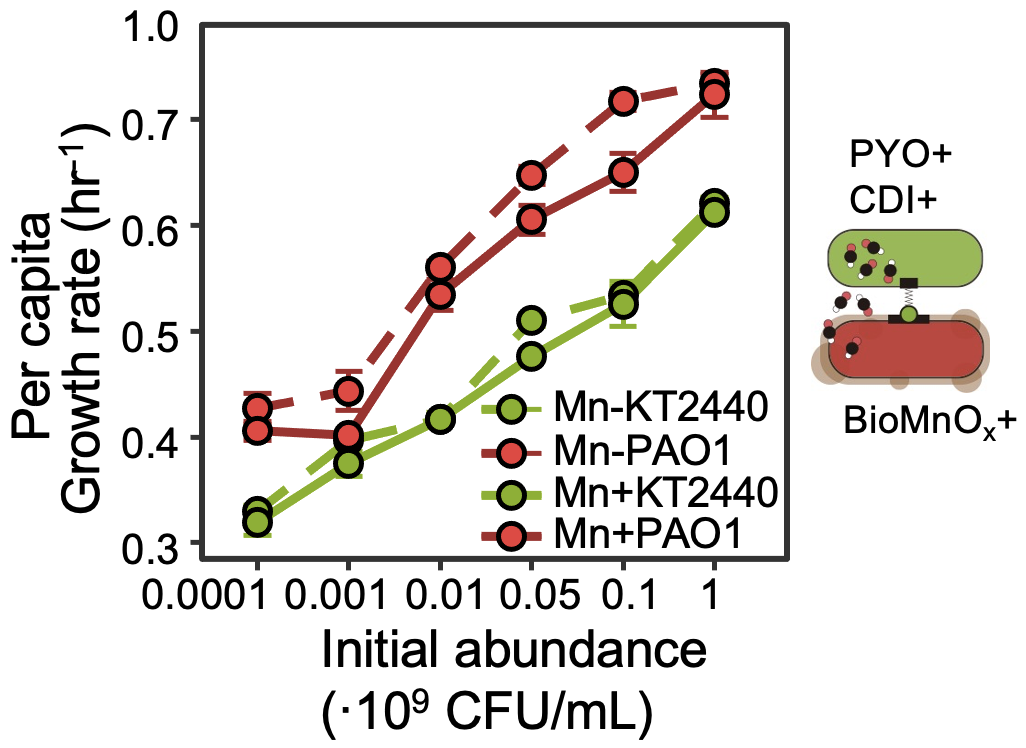

**Figure S4.** Growth rates of *P. aeruginosa* PAO1 and *P. putida* KT2440 were measured under absence and presence of 0.1 mM of MnCl_2_. Per capita growth rate refers to population growth rate normalized by initial population size (n = 6).

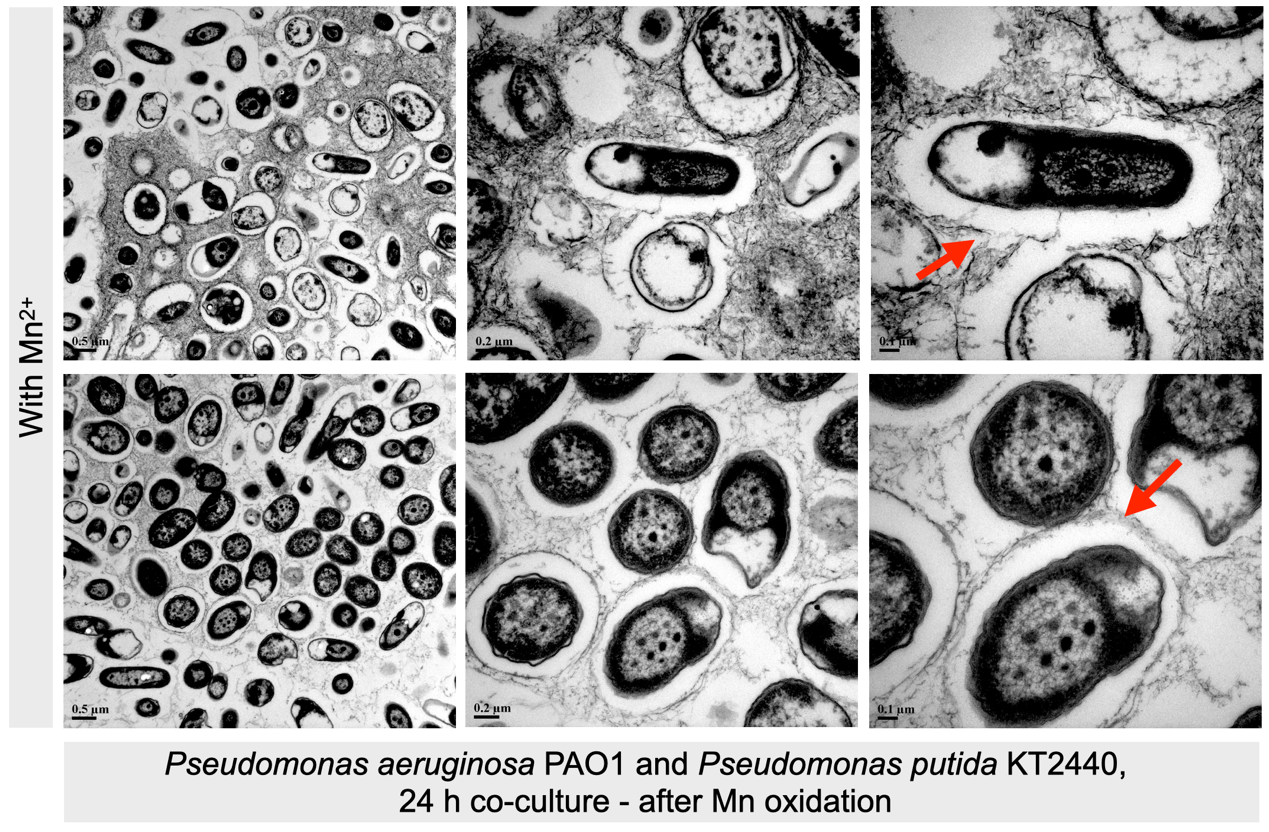

**Figure S5.** **TEM images of** P. putida **KT2440 and** P. aeruginosa **PAO1 co-culture.** TEM images after 24 h of co-culture, showing Mn(II) oxidation. Main image is marked with a scale bar of 0.5 µm, with zoomed-in details at 0.2 µm and 0.1 µm.

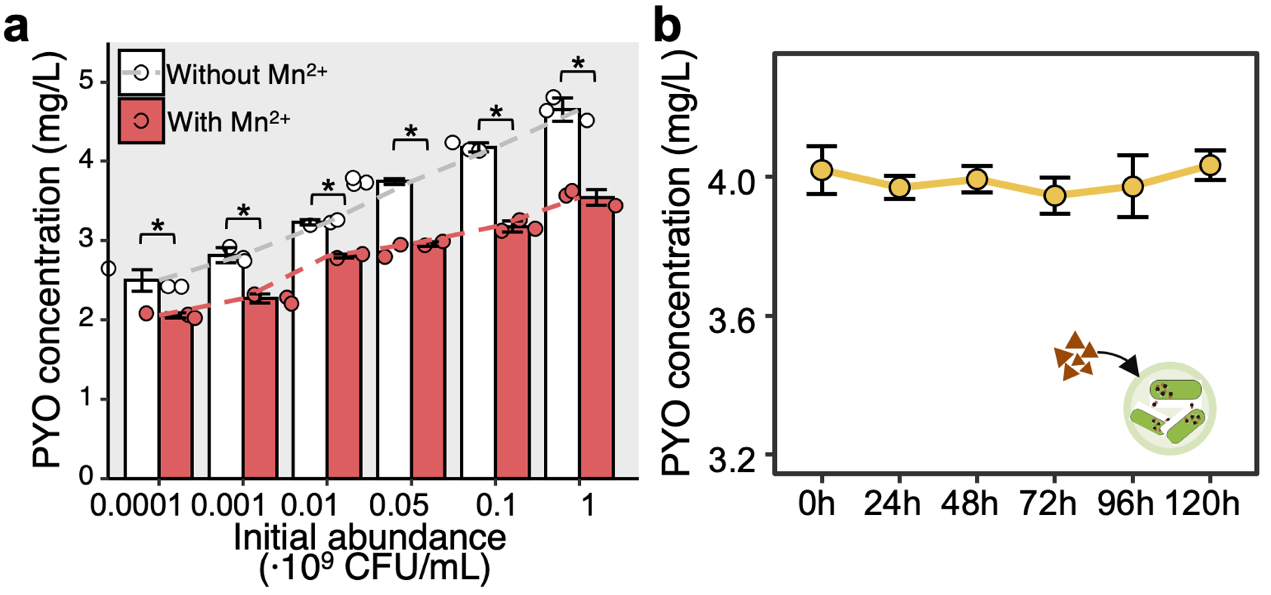

**Figure S6.** (**a**) Pyocin (PYO) production under co-culture conditions with and without MnCl₂ addition. PYO production was assessed under co-culture conditions and sterile Mn oxides were isolated from *P. putida* KT2440 monocultures. These oxides were then introduced to *P. aeruginosa* PAO1 monocultures that produced PYO. (**b**) Analysis showed no change in PYO concentration, indicating that Mn oxides did not protect against long-range weapons through the degradation of these compounds. Significance (**p* < 0.05) was assessed using Wilcoxon test (n = 6).

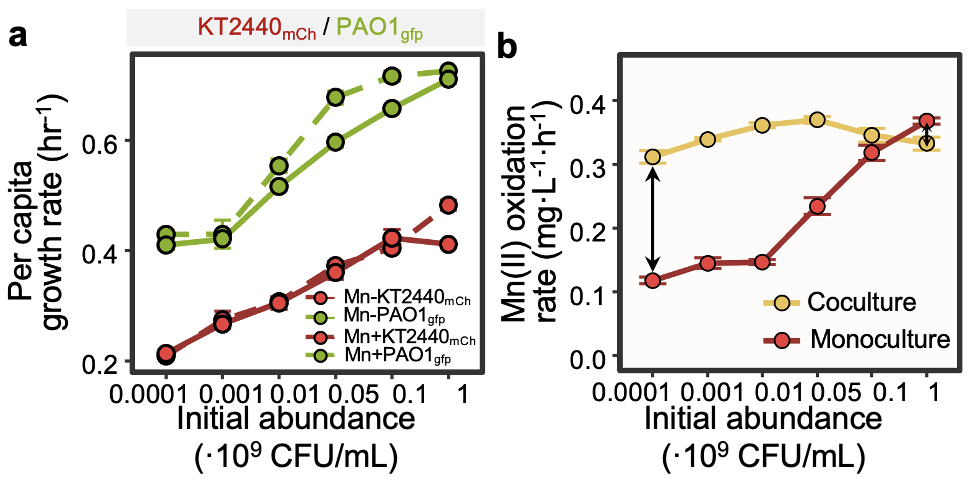

**Figure S7.** Growth and Mn(II) oxidation rates of fluorescent strains. (**a**) Growth rates of *P. aeruginosa* PAO1_gfp_ and *P. putida* KT2440_mCh_ were assessed with and without 0.1 mM MnCl₂. Per capita growth rate is population growth normalized by initial population size (n = 6). (**b**) Mn(II) oxidation rates by KT2440_mCh_, measured across various inoculum densities in co-culture with PAO1_gfp_ and in monoculture (n = 3). Significance (**p* < 0.05) was assessed using the Wilcoxon test.

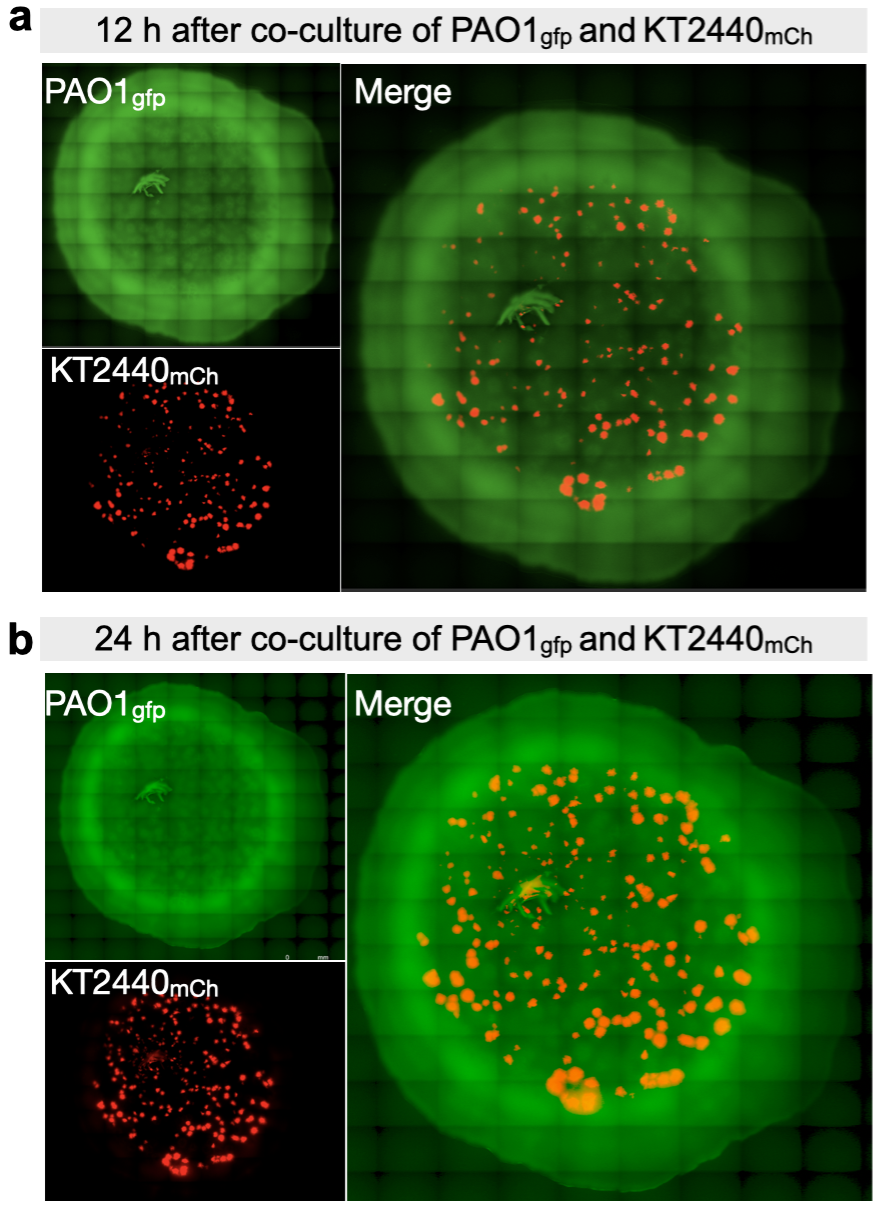

**Figure S8.** Fluorescence microscopy of *P. aeruginosa* PAO1_gfp_ and *P. putida* KT2440_mCh_ on agar plates. PAO1 (expressing GFP, green fluorescent protein) and KT2440 cells (expressing mCherry, red fluorescent protein) were co-cultured with MnCl₂ after (**a**) 12 and (**b**) 24 h of growth.

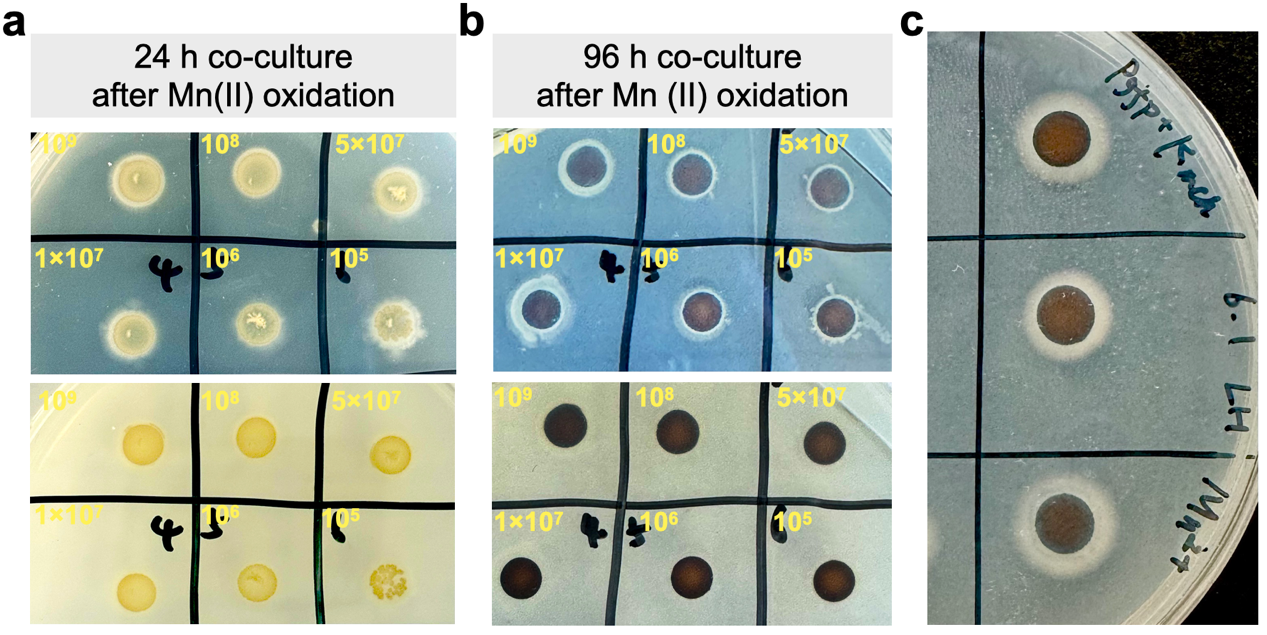

**Figure S9.** Mn(II) oxidation in co-culture of *P. aeruginosa* PAO1_gfp_ and *P. putida* KT2440_mCh_. Figures display the formation of Mn(II) oxidation on plates following co-culture with 0.1 mM MnCl_2_ (**a**) after 24 h and (**b**) after 96 h, with (**c**) a magnified view.

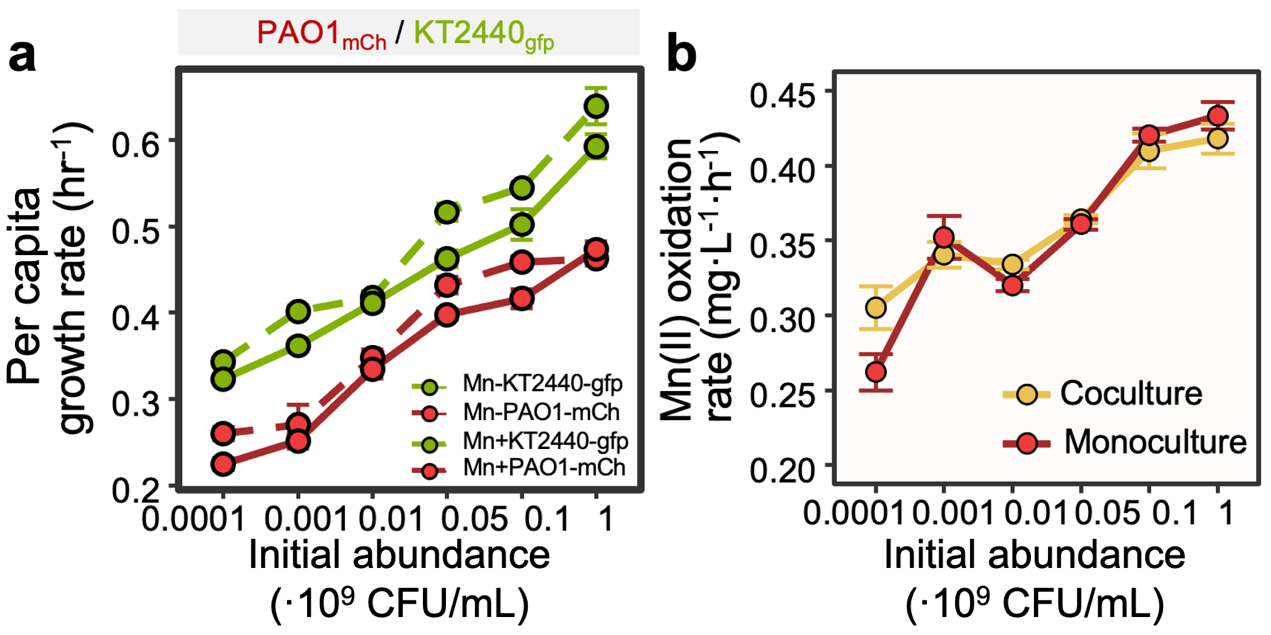

**Figure S10.** Growth and Mn(II) oxidation rates of fluorescent strains with reversed growth rate rankings. (**a**) Growth rates of *P. aeruginosa* PAO1_mCh_ and *P. putida* KT2440_gfp_ were assessed with and without 0.1 mM MnCl₂. Per capita growth rate is population growth normalized by initial population size (n = 6). (**b**) Mn(II) oxidation rates by KT2440_gfp_, were measured across various inoculum densities in co-culture with PAO1_mCh_ and in monoculture (n = 3). Significance (**p* < 0.05) was assessed using Wilcoxon test.

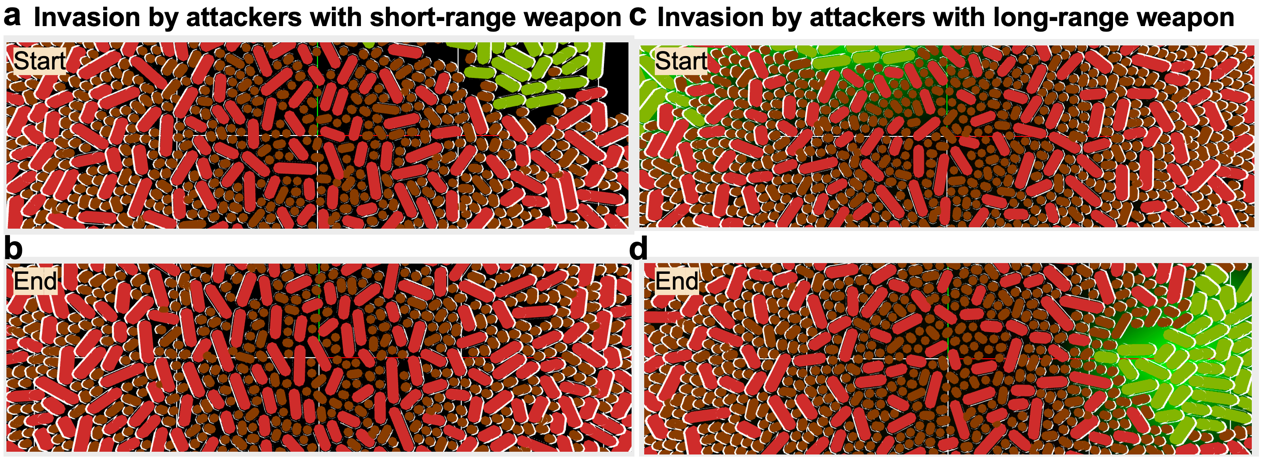

**Figure S11.** Simulation of attacker invasion in Mn(II)-oxidizing microbial communities. Subfigures illustrate the simulation dynamics of invasion where attackers are equipped with: (**a**)-(**b**) short-range weapons, demonstrating initial impacts and subsequent interactions within the communities; (**c**)-(**d**) long-range weapons, showing the initial invasion and the effectiveness of these weapons over time.

**SUPPLEMENTARY TABLES**

**Table S1** Strains and plasmids used in this study

| **Strain or plasmid** | **Relevant characteristics** | **Source** |
| --- | --- | --- |
| ***Pseudomonas aeruginosa*** |  |  |
| PAO1 | Wild-type, CDI+, PYO secretion+, SMX^R^ |  |
| KT2440 | Wild-type, CDI-, PYO secretion-, Gm^R^ |  |
| PAO1_gfp_ | Wild-type, CDI+, PYO secretion+, gfp+, Tc^R^ | This study |
| PAO1_mCh_ | Wild-type, CDI+, PYO secretion+, mCh+, Tc^R^ | This study |
| KT2440_gfp_ | Wild-type, CDI-, PYO secretion-, gfp+, Tc^R^ | This study |
| KT2440_mCh_ | Wild-type, CDI-, PYO secretion-, mCh+, Tc^R^ | This study |
| ***Escherichia coli*** | |  |
| WM3064 | Auxiliary bacteria for parental junction: *thrB1004 pro thi rpsL hsdS lacZΔM15 RP4-1360 Δ(araBAD)567 ΔdapA1341::[erm pir]* | Sangon Biotech |
| **Plasmids** | | |
| pRK415-K-EGFP::Tc | KT2440-gfp vector, Tc^R^ | Sangon Biotech |
| pRK415-K-mCherry::Tc | KT2440-mCh vector, Tc^R^ | Sangon Biotech |
| pRK415-P-EGFP::Tc | PAO1-gfp vector, Tc^R^ | Sangon Biotech |
| pRK415-P-mCherry::Tc | PAO1-mCh vector, Tc^R^ | Sangon Biotech |

**Table S2** Composition of *Pseudomonas* agar medium (PDP) used in this study

| **Chemical** | **Concentration** |
| --- | --- |
| Tryptone | 20 g/L |
| Glycerol | 10 % |
| KNO_3_ | 57.4 mM |
| MgSO_4_•7H_2_O | 14 mM |
| KH_2_PO_4_ | 50 mM |
| HEPES [4-(2-hydroxyethyl)-1-piperazineethanesulfonic acid] | 10 mM |
| Vitamin B_1_ | 0.001 mg/L |
| Vitamin B_12_ | 0.002 mg/L |
| Biotin | 0.003 mg/L |
| Agar | 15% |

**Table S3** Selective agar media used in this study

| **Species type** | **Species** | **Selective plate** | **Incubation temperature (°C)** | **Time of colony appearance (hours)** | **Colony morphology** |
| --- | --- | --- | --- | --- | --- |
| **Attacker** | *Pseudomonas aeruginosa* PAO1 | CNA (4 g/100 mL) + sulfamethoxazole (14.25 μg/mL) | 30 | 12 | Green colonies with expansive biofilms |
| **MnOB** | *Pseudomonas putida* KT2440 | CNA (4 g/100 mL) + ampicillin (10 μg/mL) | 30 | 12 | White colonies with opaque edges |
| **Attacker-gfp/mCh** | *Pseudomonas aeruginosa* PAO1 gfp/mCh | CNA (4 g/100 mL) + sulfamethoxazole (14.25 μg/mL) + tetracycline (10 μg/mL) | 30 | 12 | Green colonies |
| **MnOB-gfp/mCh** | *Pseudomonas putida* KT2440 gfp/mCh | CNA (4 g/100 mL) + ampicillin (10 μg/mL) + tetracycline (10 μg/mL) | 30 | 12 | Green colonies |

**Table S4 Model parameters used in this study**

| Type | Parameter | Symbol | Value | Unit |
| --- | --- | --- | --- | --- |
| Attackers | CDI firing rate | *k_fire_* | 100.0 | Firings cell^-1^ h^-1^ |
|  | Lethal hit threshold | *N_hits_* | 1, Infinity | - |
|  | Extracellular needle length | *L_needle_* | 0.5 | Microns |
|  | Min. needle penetration | *L_penetration_* | 0.01 | Microns |
| Cells | Max. specific growth rate | *V_max1_* | 0.45 | h^-1^ |
|  | Max. specific growth rate | *V_max2_* | 0.7 | h^-1^ |
|  | Cell radius | *R* | 0.5 | Microns |
|  | Cell volume at birth | *V_0_* | 1.16 | Microns^3^ |
|  | Cell division volume noise | *η_div_* | 9 | % |
|  | Cell division orientation noise | *η_orient_* | 0.2 | % |
| BioMnO_x_ secretion | BioMnO_x_ particle radius | *R*_BioMnOx_ | 0.4 | Microns |
|  | BioMnO_x_ particle segment length | *L*_BioMnOx_ | 0.01 | Microns |
|  | BioMnO_x_ secretion probability per time step | *P*_BioMnOx_ | 0.0-0.1 | - |
|  | BioMnO_x_ secretion cost | *c*_BioMnOx_ | 0.0 | - |
| Numerical | Simulation time step | *Δt* | 0.025 | H |
|  | Cell/needle sorting grid size | *h* | 10 | Microns |
|  | Conjugate gradient absolute tolerance | *e_CG_* | 0.001 | - |
|  | Max. contact iterations | *Max_iter_* | 8 | - |
|  | Regularization weight | *α* | 0.04 | - |
|  | Growth restriction factor | *1/γ* | 0.002 | - |
